# Intrinsic Neural Timescales in Autism Spectrum Disorder and Schizophrenia. A Replication and Direct Comparison Study

**DOI:** 10.1101/2022.06.26.497652

**Authors:** Lavinia Carmen Uscătescu, Martin Kronbichler, Sarah Said-Yürekli, Lisa Kronbichler, Vince Calhoun, Silvia Corbera, Morris Bell, Kevin Pelphrey, Godfrey Pearlson, Michal Assaf

## Abstract

Intrinsic neural timescales (INT) reflect the duration for which brain areas store information. A posterior – anterior hierarchy of increasingly longer INT has been revealed in both typically developed individuals (TD), as well as patients diagnosed with autism spectrum disorder (ASD) and schizophrenia (SZ), though INT are, overall, shorter in both patient groups. In the present study, we attempted to replicate previously reported group differences by comparing INT of TD to ASD and SZ. We replicated the previously reported result showing reduced INT in the left lateral occipital gyrus and the right post-central gyrus in SZ compared to TD. For the first time, we also directly compared the INT of the two patient groups and found that these same two areas show significantly reduced INT in SZ compared to ASD. In ASD, significant correlations were found between INT and their clinical and phenotypic characteristics. Our results point to the left lateral occipital gyrus and the right post-central gyrus as holding potential for further diagnostic refinement of SZ.

## 1. Introduction

Intrinsic neural timescales (INT) reflect the duration for which information is stored in specific brain areas (Hasson et al., 2015; Himberger et al., 2018) and are instrumental to information processing in the brain (Golesorkhi et al., 2021). In the human brain, INT length increases from posterior to anterior areas (Kiebel, Daunizeau, and Friston 2008; Shafiei et al. 2020; Raut, Snyder, and Raichle 2020). It has been suggested that caudal unimodal areas show shorter INT in order to enable the processing of slow, contextual changes (Wang 2002; Mante et al. 2013). On the other hand, the longest INT are found in anterior, higher cognitive brain areas, which perform the final integration and analysis of sensory inputs (Kiebel, Daunizeau, and Friston 2008; Wiskott and Sejnowski 2002).

Information integration requires fast and accurate information processing and transmission, but this is impaired in clinical samples. Patients diagnosed with schizophrenia (SZ) or autism spectrum disorder (ASD) show alterations in basic visual and auditory processing (e.g., Gold et al., 2012; Javitt and Freedman, 2015; Schelinski, Roswandowitz, & von Kriegstein, 2017; Schelinski & von Kriegstein, 2019; Schelinski et al., 2020), as well as motion detection (Chen et al. 2003) and object recognition (Kronbichler et al. 2018). Furthermore, multisensory integration has also been shown to be impaired in ASD and SZ. For example, ASD show reduced proneness to perceive experimentally induced audio-visual illusions such as the McGurk effect (SZ: White et al. 2014; ASD: Zhang et al. 2019), and atypical habituation to joint auditory and tactile stimulation (Green et al., 2016, 2019), as well as opposite neural activation patterns during joint visual and auditory stimulation compared to TD (Jao Keehn et al., 2017). More evidence of audio-visual integration was shown in SZ, with patients having impaired sound localization performance compared to TD (de Gelder et al., 2003). Notably, sensory integration impairments in SZ have also been shown to be heritable (Li et al., 2021). A direct relationship between sensory deficits and INT was proposed by Zilio et al. (2021), who showed that people experiencing sensory impaired states such as unresponsive wakefulness syndrome or anaesthesia also show prolonged INT.

In ASD, Watanabe, Rees, and Masuda (2019) found significantly shorter INT in the primary sensory regions (visual, sensorimotor, auditory) of adult ASD compared to TD. Similar findings in an adolescent ASD group within the same study suggest that there is a developmental component to INT patterns. Similarly, in SZ, our group (Uscătescu et al.,2021) also found decreased INT in parietal and occipital areas compared to TD, which were related to symptom severity. In addition, Wengler et al. (2020) showed that symptoms such as hallucinations and delusions are primarily related to alterations in somatosensory and auditory hierarchical INT gradients. Finally, Northoff et al. (2021) showed that INT of SZ are abnormally prolonged during self-referential processes.

In the present study, we aimed to replicate previous findings regarding the pattern of INT alterations in ASD and SZ reported by Watanabe et al. (2019) and Uscătescu et al. (2021), respectively. Both studies rely on the same computational approach to define the INT of resting-state fMRI time series. Specifically, an autocorrelation function was calculated for each voxel at incremental time lags until its value became negative. The positive autocorrelation values were then summed up and multiplied by the repetition time (TR), thus resulting in the INT index.

Using resting-state data collected from both SZ and ASD in the same site with an identical protocol, we applied a ROI analysis to focus specifically on the areas highlighted by these two studies to explore INT differences between each patient group and controls. Moreover, because of the suggested overlap between ASD and SZ in symptoms (Kästner et al., 2015) and underlying neuropathology (Moreau et al., 2021), we directly compared the two patient groups. Finally, we also performed exploratory whole-brain analyses to capture INT pattern characteristics of the three groups. To our knowledge, this is the first study to report such a comparison. Finally, we assessed the relationship between clinical and phenotypic characteristics and INT of the two patient groups.

## 2. Methods

### 2.1. Participants

Participants were recruited via the Olin Neuropsychiatry Research Center (ONRC) at the Institute of Living, Hartford Hospital, and the Department of Psychiatry, Yale School of Medicine, and underwent resting-state fMRI scanning for the current study. After discarding datasets displaying head motion > 10 mm, our dataset contained 58 TD, 39 ASD, and 41 SZ. Of these, some were subsequently excluded due to incomplete phenotypic assessment information, thus resulting in the following final samples: 55 TD, 30 ASD, and 39 SZ. As this dataset has been previously used by Hyatt et al. (2020, 2021) and Rabany et al. (2019), the exclusion criteria were the same, namely: intellectual disability (i.e., estimated IQ < 80), a neurological disorder (e.g., epilepsy), current drug use as indicated by pre-scanning interview and urine test, incompatibility with MRI safety measures (e.g., ferromagnetic implants), and a history of psychiatric and neurological diagnoses in TD.

### 2.2. Clinical and phenotypical assessment

The severity of psychotic symptoms was assessed using the Positive and Negative Syndrome Scale (PANSS; Kay et al., 1987) in both the ASD and SZ groups. The PANSS scores can be interpreted along three subscales: positive symptoms, reflecting the severity of hallucinations and delusions; negative symptoms, reflecting the severity of blunted affect and anhedonia, and a general subscale quantifying other psychopathologies such as poor attention and lack of insight. The ADOS, module 4 (Lord et al., 2000) was administered to all participants to confirm/rule out an ASD diagnosis. ADOS total score (social interaction and communication subscores) was used to measure autism-related symptom severity. The Intelligence Quotient (IQ) was estimated for the entire sample using the Vocabulary and Block Design subtests of the Wechsler Scale of Adult Intelligence-III (WAIS-III; Wechsler, 1997; Sattler and Ryan, 1999). The structured clinical interview for DSM-IV-TR axis I disorders (SCID; First et al., 2002) was additionally used to confirm SZ diagnosis and the absence of any Axis I diagnoses in TD. To assess different social cognitive abilities, all participants were asked to complete the Empathy Quotient (EQ; Wakabayashi, Baron-Cohen & Wheelwright, 2006) and the Bermond–Vorst Alexithymia Questionnaire (BVAQ; Vorst & Bermond, 2001). BVAQ subscores are computed along five distinct dimensions: “verbalizing” reflects one’s propensity to talk about one’s feelings; “identifying” reflects the extent to which one can accurately define one’s emotional states; “analyzing” quantifies the extent to which one seeks to understand the reason for one’s emotions; “fantasizing” quantifies one’s tendency to day-dream, and “emotionalizing” reflects the extent to which a person is emotionally aroused by emoting inducing events. Means and standard deviations, as well as group comparison tests of the above-mentioned instruments, are given in Table 1.

**Table 1.** Means and standard deviations (in parentheses) of demographics, phenotypic and clinical instrument scores for all three groups. FD = framewise displacement; est. IQ = estimated Intelligence Quotient; EQ = Empathy Quotient; ADOS = Autism Diagnostic Observation Schedule module 4; BVAQ = Bermond–Vorst Alexithymia Questionnaire. Group statistics are shown in the last four columns. Pairwise comparisons were performed using Welch’s two-sample t-test. Both uncorrected (i.e., *p*) and false discovery rate corrected (i.e., *p*_FDR_) *p* values are shown.

|  | TD | ASD | SZ | ASD v. SZ v. TD | ASD > TD | ASD > SZ | TD > SZ |
| --- | --- | --- | --- | --- | --- | --- | --- |
| Females/Males | 29/26 | 5/25 | 8/31 | $\chi^2(2) = 15.8$,<br>.<.000 | | | |
|  |  |  |  | **F(2,121), $p$** | **t(df), $p$, $p_{\text{FDR}}$** | **t(df), $p$, $p_{\text{FDR}}$** | **t(df), $p$, $p_{\text{FDR}}$** |
| Age (year) | 24<br>(3.73) | 22<br>(3.74) | 26<br>(3.58) |  | -1.98 (79.8),<br>.051, .051 | **-4.06 (76.9),**<br>**<.000, <.000** | **-2.46 (88), .016,**<br>**.024** |
| FD (mm) | 0.08<br>(0.03) | 0.09<br>(0.04) | 0.11<br>(0.1) | **5.23, .007** | 2.52 (66.6),<br>.014, .021 | **-0.772 (69.6),**<br>**.443, .443** | **-2.71 (54.3),**<br>**.009, .021** |
| est. IQ | 112.26<br>(14.62) | 109.1<br>(15.21) | 99.41<br>(13.34) | **9.366, <.000** | -1.33 (82.8),<br>.186, .186 | **3.26 (77), .002,**<br>**.003** | **4.97 (90.5),**<br>**<.000, <.000** |
| EQ | 49<br>(10.28) | 33.57<br>(11.1) | 39.8<br>(12.26) | **22.37, <.000** | **-6.73 (72.2),**<br>**<.000, <.000** | **-2.50 (73.9),**<br>**.015, .015** | **3.65 (72.2),**<br>**<.000, <.000** |
| ADOS | 1.87<br>(1.45) | 10.1<br>(2.61) | 8.41<br>(5.26) | **78.26, <.000** | **17.5 (52.2),**<br>**<.000, <.000** | 1.67 (58.8),<br>1.72, 1.72 | **-7.62 (44.2),**<br>**<.000, <.000** |
| BVAQ<br>Verbalizing | 18.55<br>(5.51) | 22.2<br>(4.54) | 22<br>(5.03) | **7.02, .001** | **3.07 (81.4),**<br>**.003, .005** | -0.0571 (75.1),<br>.955, .955 | **-3.22 (90.4),**<br>**.002, .005** |
| BVAQ<br>Fantasizing | 19.69<br>(5.12) | 17.9<br>(6.03) | 21.36<br>(5.48) | **4, .02** | -1.17 (73.5),<br>.247, .247 | **-2.72 (75.2),**<br>**.008, .024** | -1.88 (82.8),<br>.064, .096 |
| BVAQ<br>Identifying | 15.53<br>(4.82) | 18.57<br>(6.4) | 20.41<br>(4.96) | **10.43, <.000** | **2.59 (63), .012,**<br>**.018** | -1.29 (68.2),<br>.203, .203 | **-4.79 (85.2),**<br>**<.000, <.000** |
| BVAQ<br>Emotionalizing | 22.38<br>(3.74) | 21.83<br>(3.51) | 22.56<br>(4.2) | 0.6, .55 | -0.810 (81.4),<br>.421, .632 | -1.07 (74), .289,<br>.632 | -0.389 (77.9),<br>.698, .698 |
| BVAQ<br>Analyzing | 17.91<br>(4.43) | 18.47<br>(4.27) | 19.9<br>(4.5) | 2.81, 0.6 | 0.576 (81.9),<br>.566, .566 | -1.62 (76), .11,<br>.165 | -2.27 (85.4),<br>.026, .078 |
| PANSS<br>Positive |  | 12.1<br>(2.86) | 15.36<br>(4.86) |  |  | **-3.88 (66.96),**<br>**<.000** |  |
| PANSS<br>Negative |  | 15.57<br>(4.7) | 19.26<br>(6.2) |  |  | **-2.644 (72.75),**<br>**.01** |  |
| PANSS<br>General |  | 26.7<br>(5.62) | 31.59<br>(6.98) |  |  | **-3.716 (73),**<br>**<.000** |  |

### 2.3. Imaging data acquisition, preprocessing and motion correction

All resting-state fMRI scans lasted 7.5 min and were collected using a Siemens Skyra 3 T scanner at the ONRC. Participants lay still, with eyes open, while fixating on a centrally presented cross. Blood oxygenation level-dependent (BOLD) signal was obtained with a T2*-weighted echo-planar (EPI) sequence: R/TE = 475/30 msec, flip-angle = 60°, 48 slices, multiband (8), interleaved slice order, 3 mm^3^ voxels.

Pre-processing of structural MRI data was done using fMRIPrep v20.2.6 (Esteban et al., 2018), which is based on Nipype 1.7.0 (Gorgolewski et al., 2011; Gorgolewski et al., 2018). The pipeline included the following steps: correction for intensity non-uniformities using field maps (Tustison et al. 2010), skull-stripping using ANTs 2.3.3 (Avants et al. 2008) with OASIS30ANTs as target template, segmentation using the FAST algorithm from FSL 5.0.9 (Zhang, Brady, and Smith 2001), and normalization using antsRegistration (ANTs 2.3.3). Functional scans were co-registered with FLIRT (FSL 5.0.9, Jenkinson and Smith 2001) using nine degrees of freedom and spatiotemporal filtering was performed using MCFLIRT (FSL 5.0.9, Jenkinson et al. 2002). Finally, a slice-time correction was applied (Parker & Razlighi, 2019).

Motion artifacts were first removed using non-aggressive ICA-AROMA (Pruim et al., 2015), following smoothing with a 6mm FWHM kernel. Detrending was then performed using DiCER (Aquino et al., 2020). Finally, frame-wise displacement (FD) motion parameters were computed according to the FSL library algorithm (Jenkinson et al., 2012) and later used as covariates to check that our results were not biased by potential motion artifacts.

### 2.4. Data analysis

The INT analysis steps described in Watanabe, Rees and Masuda (2019) and implemented in Uscatescu et al. (2021) were followed. First, an autocorrelation function (ACF) was calculated for each voxel. The ACF measures how data points in a time series are related to each other, or in other words, the self-similarity of the rsfMRI BOLD signal. First, we set a maximum time lag of 20 seconds and divide it into smaller, incremental timesteps/time lags for each second. At each timestep, we correlate the preceding and the current signal and proceed thus until the value of the correlation turns negative. Finally, the resulting positive autocorrelation values at each voxel were summed up, and this value was then multiplied by the TR, thus resulting in the final INT index.

Whole-brain analyses were performed using the SPM12 software (http://www.fil.ion.ucl.ac.uk/spm/) while the regions of interest (ROIs) were defined using the MARSBAR toolbox (Brett et al., 2002). Further statistical analyses were performed in R 5.263 software.

The ROIs were defined as 6 mm radius spheres centered around the peak MNI coordinates reported by Watanabe et al. (2019) and Uscătescu et al. (2021). The peak coordinates of all 13 ROIs are shown in Table 2 (note that regions derived from Watanabe et al. (2019) are prefixed with ‘W’ and those from Uscătescu et al. (2021) with ‘U’).

**Table 2.** Peak MNI coordinates reported by Watanabe et al. (2019) and Uscătescu et al. (2021), around which the ROIs of the current study were defined.

| ROI |  | MNI Coordinates |  |  |
| --- | --- | --- | --- | --- |
| ROI Acronym | ROI Description | X | Y | Z |
| Watanabe et al. (2019) |  |  |  |  |
| W_rPCG | right Post-Central Gyrus | 58 | -14 | 44 |
| W_lPCG | left Post-Central Gyrus | -58 | -14 | 40 |
| W_rMTG | right Middle Temporal Gyrus | 60 | 2 | 26 |
| W_lMTG | left Middle Temporal Gyrus | -70 | -26 | 6 |
| W_rIOG | right Inferior Occipital Gyrus | 52 | -74 | 6 |
| W_rIPL | right Inferior Parietal Lobule | 50 | -44 | 32 |
| W_rMidIns | right Middle Insula | 50 | 10 | 4 |
| W_rCaud | right Caudate | 14 | 20 | 12 |
| Uscătescu et al. (2021) |  |  |  |  |
| U_rOccFusG | right Occipital Fusiform Gyrus | 27 | -88 | 14 |
| U_lSupOccG | left Superior Occipital Gyrus | -15 | -91 | 19 |
| U_rSupOccG | right Superior Occipital Gyrus | 18 | -79 | 25 |
| U_lLatOccC | left Lateral Occipital Cortex | -42 | -70 | 1 |
| U_rPostCenG | right Post-Central Gyrus | 42 | -31 | 64 |

## 3. Results

### 3.1. Group differences in clinical and phenotypical assessment

Age (i.e., ASD < TD < SZ), estimated IQ (i.e., SZ < ASD < TD) and sex (i.e., more males than females) were significantly different between groups (see Table 1). Therefore, these variables (as well as FD, as described above) were used as covariates in group-comparison analyses.

### 3.2. ROI replication results

First, we explored overall group differences by running an ANCOVA analysis with age, sex, IQ, and FD as covariates (summarized in Table 3). Of the ROIs based on Watanabe et al. (2019), only the right middle insula (W_rMidIns) showed a significant main effect (F (2, 131) = 3.31, *p* = .04, ηp2 = 0.05), but not after false discovery rate (FDR) correction (*p*_FDR_ = .1). Of the ROIs based on Uscătescu et al. (2021), four out of five showed a significant main effect that also survived FDR correction (see Table 3).

**Table 3.** Group differences in INT per ROI, calculated with ANCOVA with age, sex, IQ, and FD as covariates. The effect size was calculated using partial eta squared (ηp2). False discovery rate *p* values (***p***_***FDR***_) are also shown. Results in bold font reflect post-FDR significant comparisons.

| | | INT mean (sd) | | | F(2,131) | $p$ | $p_{FDR}$ | $\eta^2$ |
| --- | --- | --- | --- | --- | --- | --- | --- | --- |
|  |  | TD | ASD | SZ |  |  |  |  |
| 1 | W_rPCG | 2.88<br>(1.36) | 2.8<br>(1.16) | 2.37<br>(1.21) | 2.74 | .07 | .11 | 0.04 |
| 2 | W_IPCG | 2.83<br>(1.27) | 2.7<br>(1.08) | 2.44<br>(1.2) | 2.85 | .06 | .11 | 0.04 |
| 3 | W_rMTG | 1.3<br>(0.8) | 1.46<br>(0.99) | 1.21<br>(1.22) | 0.38 | .69 | .69 | 0.01 |
| 4 | W_IMTG | 1.85<br>(0.95) | 2.07<br>(1.06) | 1.66<br>(1.18) | 1.49 | .23 | .3 | 0.02 |
| 5 | W_rIOG | 2.16<br>(1.12) | 2.19<br>(1.39) | 1.67<br>(1.18) | 2.28 | .11 | .16 | 0.03 |
| 6 | W_rIPL | 2.15<br>(0.93) | 2.25<br>(1.17) | 2<br>(1.18) | 0.46 | .63 | .68 | 0.01 |
| 7 | W_rMidIns | 2.12<br>(0.93) | 1.98<br>(1.11) | 1.65<br>(1.16) | 3.31 | .04 | .1 | 0.05 |
| 8 | W_rCaud | 0.79<br>(0.67) | 0.88<br>(0.95) | 0.76<br>(1.06) | 0.78 | .46 | .54 | 0.01 |
| 9 | **U_rOccFusG** | 2.57<br>(1.28) | 2.37<br>(1.22) | 2.2<br>(1.22) | **5.65** | **.004** | **.03** | **0.08** |
| 10 | U_lSupOccG | 2.18<br>(1.2) | 2.17<br>(1.2) | 1.86<br>(1.13) | 2.8 | .07 | .11 | 0.04 |
| 11 | U_rSupOccG | **1.64**<br>(1.1) | **1.4**<br>(0.94) | **1.28**<br>(1.17) | **4.48** | **.01** | **.03** | **0.06** |
| 12 | U_lLatOccC | **2.48**<br>(1.15) | **2.26**<br>(1.1) | **1.73**<br>(1.19) | **4.78** | **.01** | **.03** | **0.07** |
| 13 | U_rPostCenG | **2.7**<br>(1.45) | **2.44**<br>(1.44) | **1.92**<br>(1.36) | **7.44** | **<.000** | **0** | **0.1** |

Further group-wise comparisons via Welch’s one-tailed, two-sample *t-*tests verified whether the group differences previously reported by Watanabe et al. (2019) and Uscătescu et al. (2021), comparing each clinical group to TD could be replicated in our current sample. We also directly compared the two patient groups (these results are summarised in Table 4). None of the ROIs based on Watanabe et al. (2019) showed significant INT group differences between TD and ASD. The right inferior occipital gyrus (W_rIOG) and the right middle insula (W_rMidIns) showed significantly increased INT in ASD compared to SZ, but did not survive FDR correction (see Table 4). Replicable group differences were found for two of the ROIs from Uscătescu et al., namely, both the left lateral occipital cortex (U_lLatOccC) and the right post-central gyrus (U_rPostCenG) displayed significantly reduced INT in SZ compared to TD (see Table 4). These two ROIs also showed significantly increased INT in ASD compared to SZ, but only before FDR correction (see Table 4).

**Table 4.**
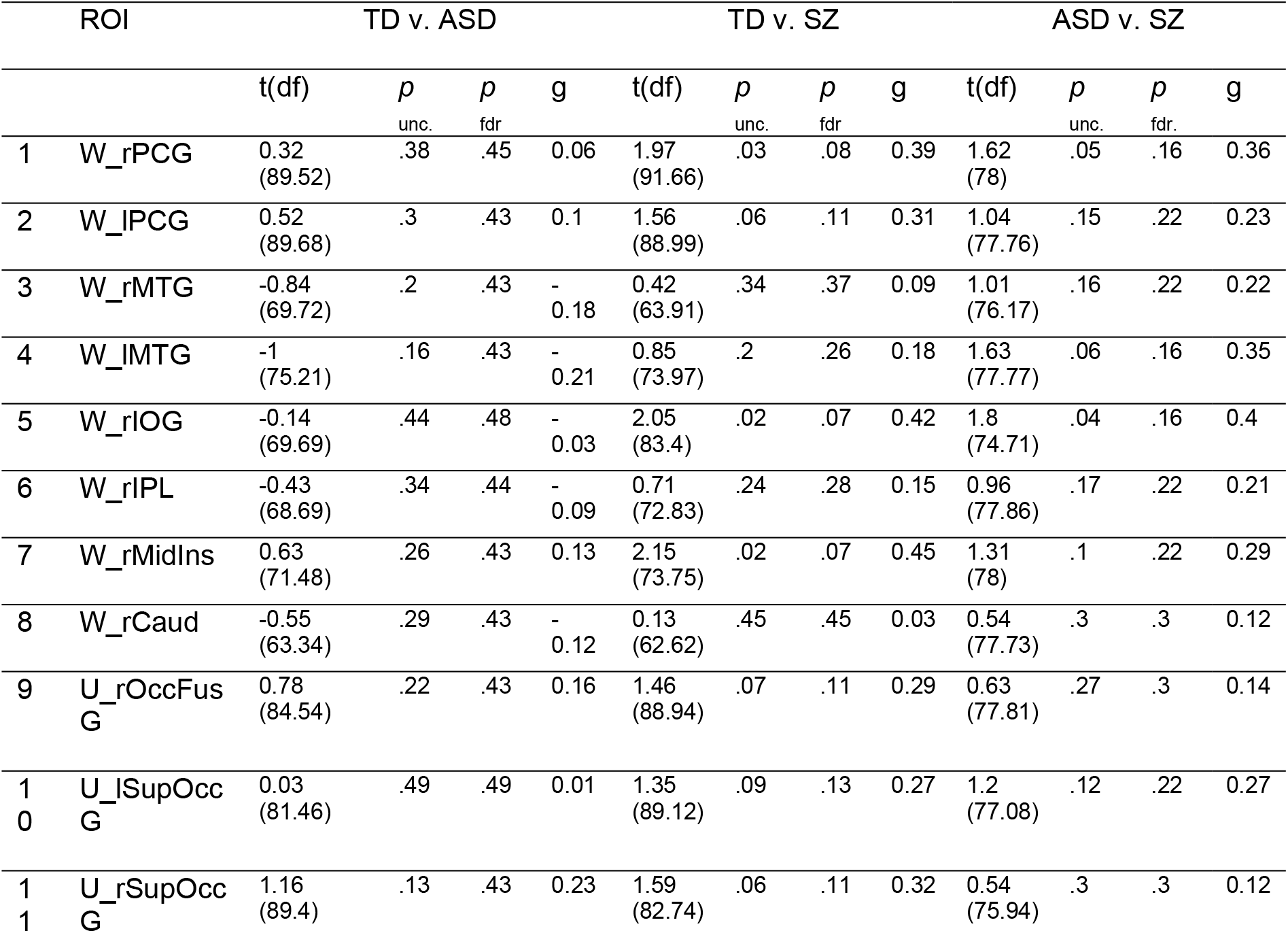
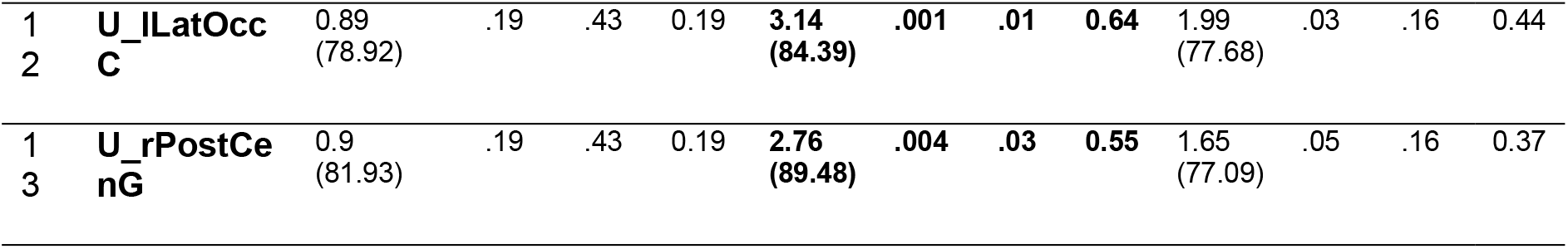
One tailed, Welch’s two-sample t-tests of INT per ROI, with and without FDR correction, with Hedge’s g effect size. Results in bold font reflect post-FDR significant comparisons.

Finally, we also assessed the relationship between clinical and phenotypic measures and INT in ASD and SZ. The significant correlations before and after FDR correction are shown in Figure 1. None of the initially significant correlations in SZ between INT and PANSS (Uscătescu et al., 2021) survived FDR correction. In the ASD sample, we replicated the negative correlations reported by Watanabe et al. (2019) between the ADOS total score and the INT of the W_rPCG, W_lPCG and W_rIOG.

**Figure 1.**
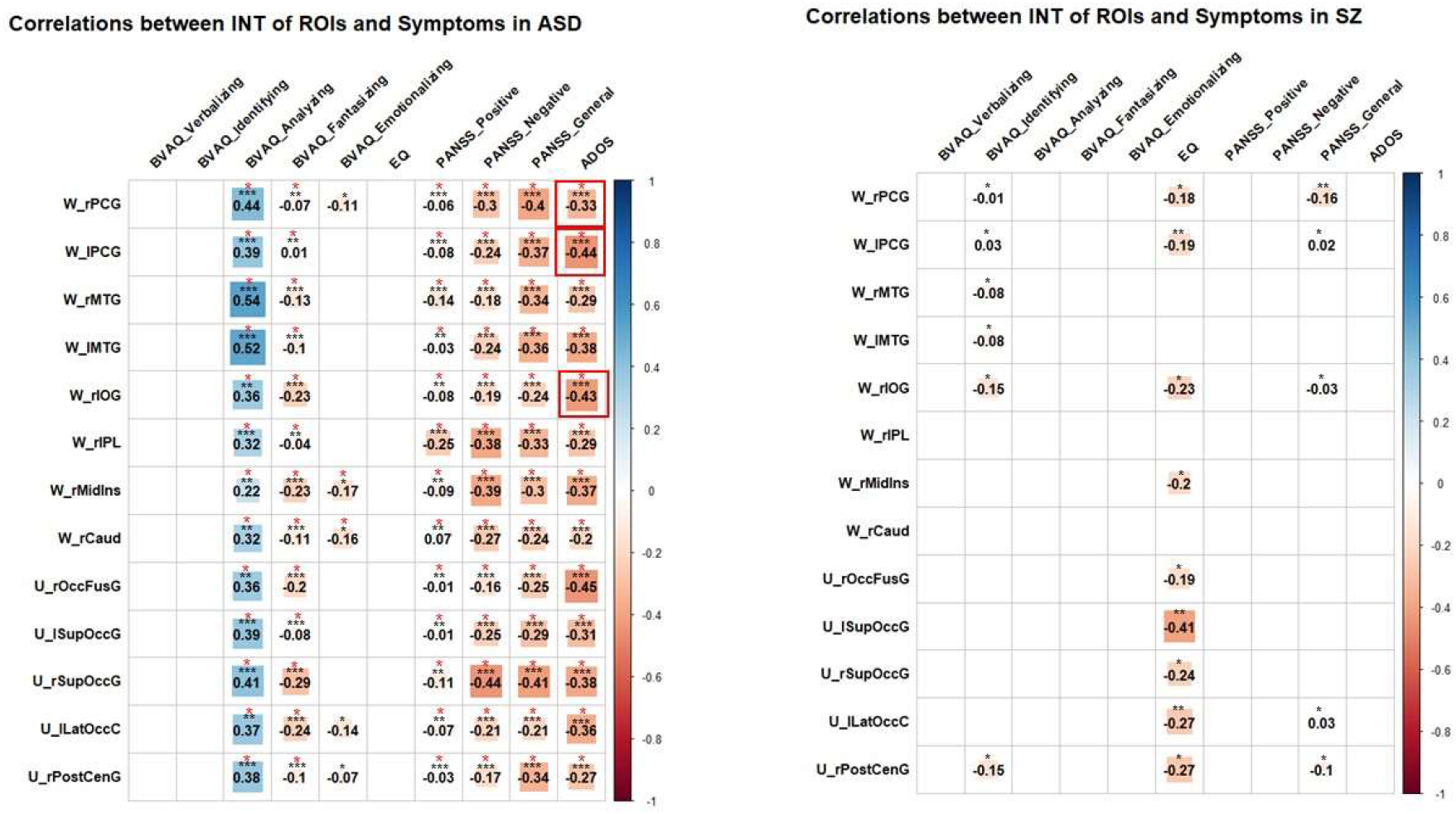
Pearson correlations between INT of ROIs and clinical measures in ASD (left) and SZ (right). Significant correlation values at p < .05, both prior to (black asterisks) and after (red asterisks) FDR, are plotted, while non-significant correlations have been replaced by blank spaces. Replicated correlations are marked with red squares.

### 3.3. Whole-brain exploratory results

In a first step, we performed mass-univariate analyses to compare brain-wise group differences in INT. The areas that exhibited INT differences across the three groups (Figure 2.A.) were the left lateral occipital gyrus (lLOG), the left supramarginal gyrus (lSMG), the right precentral gyrus (rPG), the right fusiform gyrus (rFusG) and the right inferior temporal gyrus (rITG). Within-group INT are displayed in Figure 2.B. Pairwise comparisons showed no areas displaying higher INT in TD compared to ASD, but higher INT values were revealed in ASD compared to TD (Figure 2.C bottom) in the left Fusiform Gyrus (lFusG), rITG, and right Entorhinal Cortex (rEnt). Higher INT were found in TD compared to SZ (Figure 2.C top right) in left Inferior Occipital Gyrus (lIOG), left Superior Occipital Gyrus (lSOG), left Superior Parietal Lobe (lSPL), left Pre-Central Gyrus (lPreCenG), right Superior Frontal Gyrus (rSFG), right Pre-Central Gyrus (rPreCenG), right Superior Parietal Lobe (rSPL), right Post-Central Gyrus (rPostCenG), rFusG, right Medial Temporal Gyrus (rMTG), and right Superior Frontal Gyrus (rSFG). No INT were found to be higher in SZ compared to TD. No areas displayed larger INT in SZ compared to ASD, but larger INT were found in ASD than in SZ (Figure 2.C top left) in left Medial Frontal Gyrus (lMFG), rPreCenG, rSPL, right Supra-Marginal Gyrus (rSMG), and rITG.

**Figure 2.**
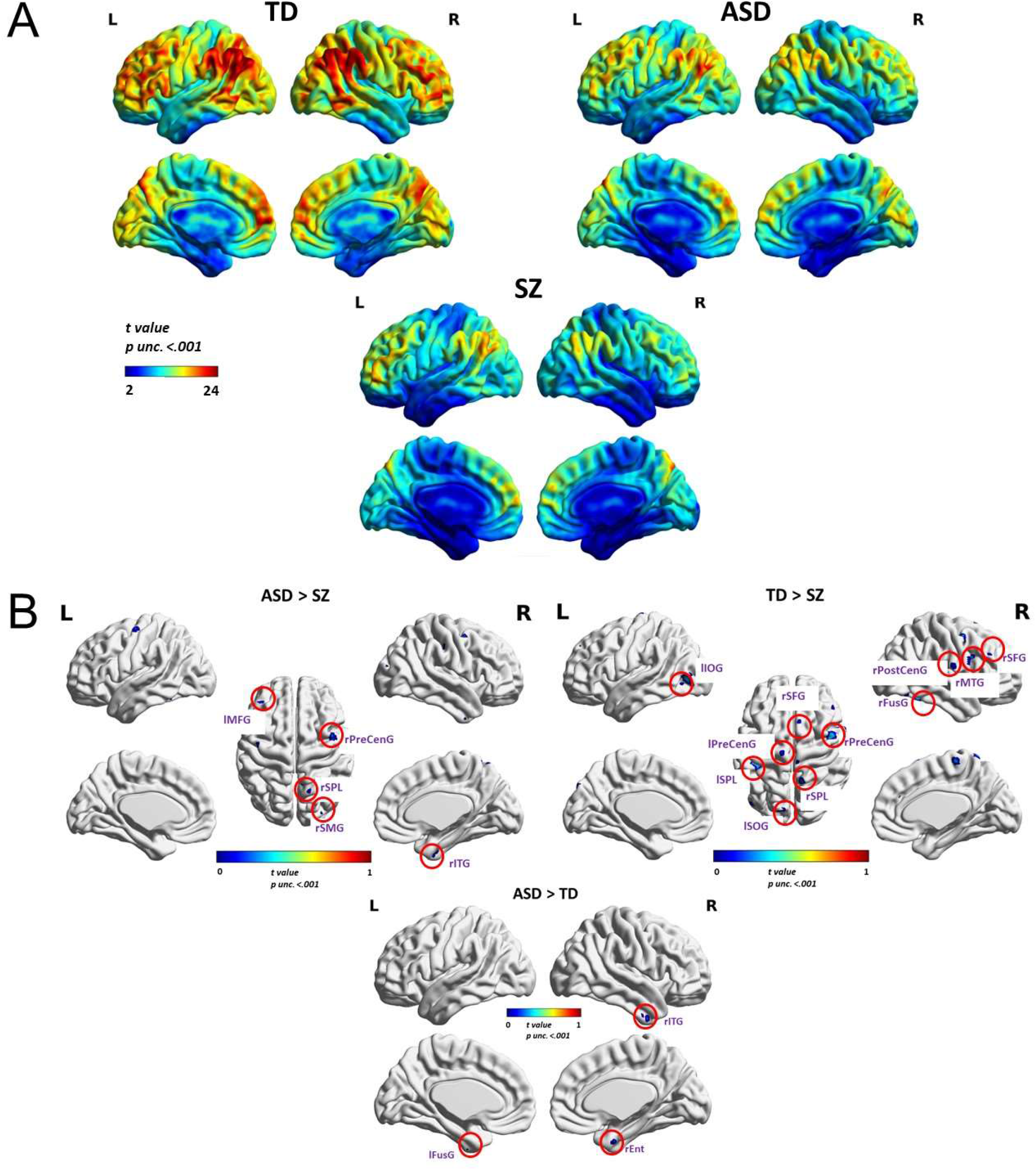
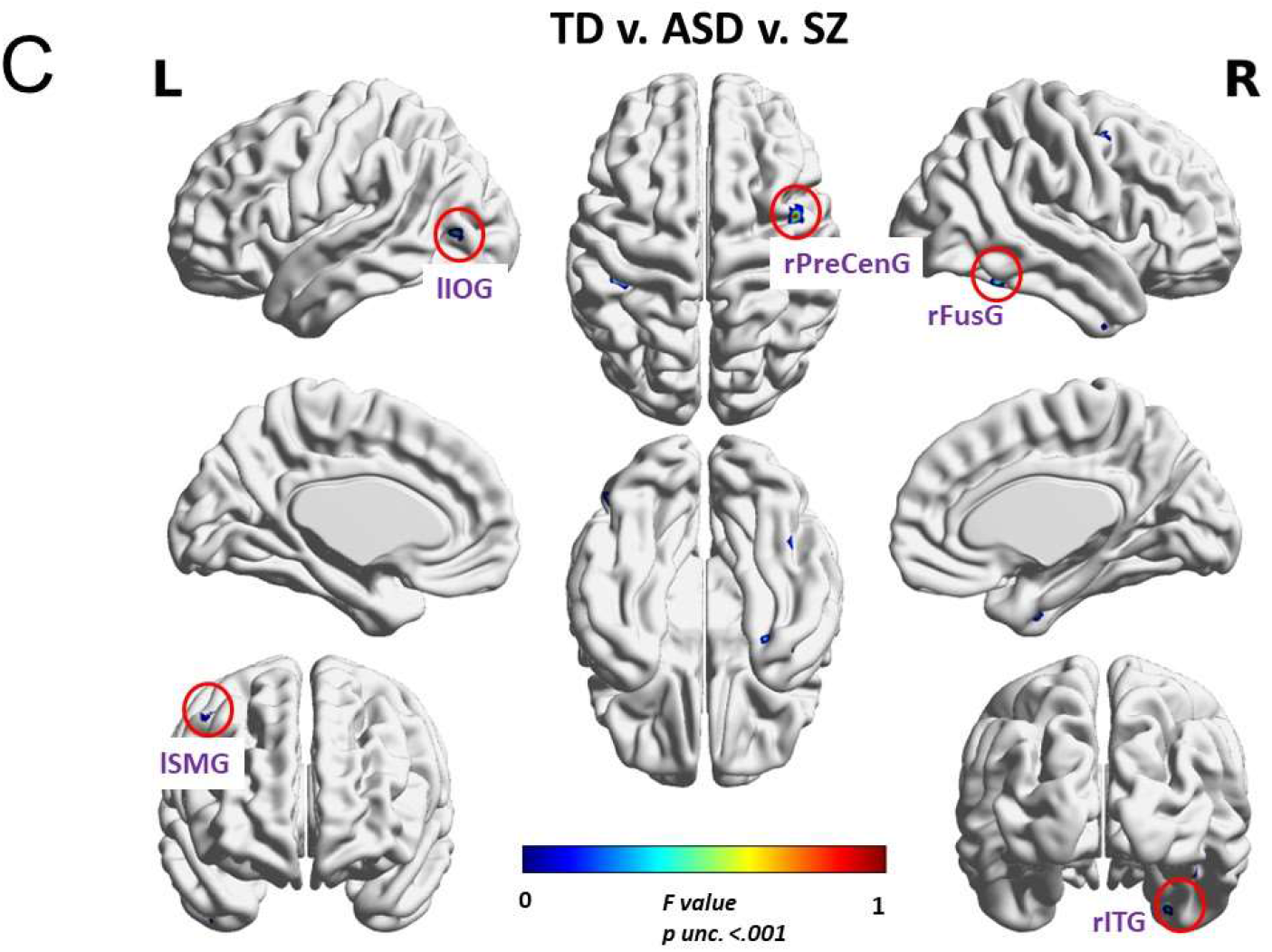
**A**. Voxel-wise INT values within each group; TD: typically developed; ASD: autism spectrum disorder; SZ: schizophrenia. **B. Top left:** Areas displaying higher INT in ASD compared to TD. **Top right:** Areas displaying higher INT in TD compared to SZ. **Bottom:** Areas displaying higher INT in ASD compared to SZ. **A**. Brain areas exhibiting INT differences across the three groups in a whole-brain ANOVA; lLOG: left Lateral Occipital Gyrus; lSMG: left Supra-Marginal Gyrus; rPreCenG: the right pre-central gyrus; rFusG: the right fusiform gyrus; rITG: the right inferior temporal gyrus; lFusG: left fusiform gyrus; rEnt: right entorhinal cortex; lMFG: left medial frontal gyrus; lPreCenG: left pre-central gyrus; rSPL: right superior parietal lobe; rSMG: right supra-marginal gyrus; rITG: right inferior temporal gyrus; lIOG: left inferior occipital gyrus; lSOG: left superior occipital gyrus; lSPL: left superior parietal lobe; rSFG: right superior frontal gyrus; rMTG: right medial temporal gyrus; rSFG: right superior frontal gyrus.

## 4. Discussion

In the current study, we sought to replicate previous findings regarding INT differences between TD and ASD, and TD and SZ independently. We also verified to which extent previously reported relationships between INT and clinical and phenotypic characteristics could be replicated. In addition, we made the novel contribution of directly comparing ASD to SZ.

Uscătescu et al. (2021) previously reported five ROIs that showed a trend of decreased INT in SZ compared to TD (right occipital fusiform gyrus/U_rOccFusG, left superior occipital gyrus/U_lSupOccG, right superior occipital gyrus/U_rSupOccG, left lateral occipital cortex/U_lLatOccC, right post-central gyrus/U_rPostCenG), two of which were replicated across three independent datasets, namely the U_rSupOccG and the U_rPostCenG. In the present study, we found a similar pattern of decreased INT in SZ compared to TD in all five ROIs, but this difference was significant only for U_rSupOccG and the U_rPostCenG. These findings suggest that the right superior occipital gyrus and the right post-central gyrus are the most reliable areas in terms of INT replicability in SZ. The right post-central gyrus has also been previously shown to exhibit increased temporal variability in SZ compared to TD (Zhang, Guo & Tian, 2019), and to be hyper-connected to the thalamus in both ASD (Fu et al., 2019) and SZ (Ferri et al., 2018) compared to TD. Increased temporal irregularities have also been previously reported in the right superior occipital gyrus in SZ compared to TD (Xue et al., 2019). Taken together, this evidence suggests that these two areas, from which INT group differences could be replicated in our study, reliably show temporal abnormalities in SZ.

We were unable to replicate the ASD v. TD group differences reported by Watanabe et al. (2019) in any of the eight ROIs from their study (right post-central gyrus/W_rPCG, left post-central gyrus/W_lPCG, right middle temporal gyrus/W_rMTG, left middle temporal gyrus/W_lMTG, right inferior occipital gyrus/W_rIOG, right inferior parietal lobule/W_rIPL, right middle insula/W_rMidIns, right caudate/W_rCaud). However, the trends that we observed with respect to the INT group differences were similar to Watanabe et al. (2019), who reported that ASD had longer INT compared to TD in the W_rCaud, and shorter ones in the bilateral postcentral gyri.

Significant associations between symptom severity and phenotypic assessment were initially found in the SZ group, namely: significant negative correlations between the INT of the W_lPCG and BVAQ Identifying, and PANSS general psychopathology, and between the INT of the U_lLatOccC and PANSS general psychopathology. Additionally, in SZ, we found a series of negative correlations between BVAQ Identifying and the INT of U_rPostCenG, W_rPCG, W_lPCG, W_rMTG, W_lMTG, W_rIOG, W_rIPL, W_rMidIns, W_rCaud, the EQ and the INT of U_rOccFusG, U_lSupOccG, U_rSupOccG, U_lLatOccC, U_rPostCenG, W_rPCG, W_lPCG, W_rIOG, W_rMidIns and between PANSS general psychopathology and the INT of W_rPCG, W_lPCG, W_rIOG, U_lLatOccC, and U_rPostCenG. However none of these correlations survived FDR correction, nor did they replicate those previously reported by Uscătescu et al. (2021). With respect to the ASD group, the significant negative correlations between the ADOS total score and the INT of W_lPCG, W_rPCG, and the W_rIOG reported before by Watanabe et al. (2019) were replicated and survived FDR correction. These brain-psychopathology correlation results should be taken cautiously, as two recent studies drew attention to how (even when large samples are available) such effects tend to be very small and rather difficult to replicate (Gratton, Nelson & Gordon, 2022; Marek et al., 2022).

Finally, we directly compared the two patient groups with respect to the INT of the 13 ROIs. In all cases, the INT of ASD were longer than those of SZ, though only significant in W_rIOG, W_rPCG, U_lLatOccC and U_rPostCenG and only before FDR correction. We believe that these results are a relevant step towards localising the sites that could hold potential for diagnostic refinement of ASD and SZ for the temporal irregularities of their neural functioning. However, given the diagnostic heterogeneity of both ASD and SZ, it is necessary to further ascertain the replicability of these findings before more decisive conclusions can be formulated. The present study has the distinct advantage that rsfMRI data from ASD and SZ and controls were recorded in the same setting, thus eliminating potential confounds related to variation in data collection. However, some limitations remain. On the one hand, some sensory areas, like the postcentral gyrus, have been shown to display decreased sensory sensitivity with age (Diaz & Yalcinbas, 2021). Although we included age as covariate in our group analysis, age-balanced samples would have been optimal. A second limitation, which also determined our choice of not eliminating participants despite the age imbalance, is related to the relatively small sample sizes. The limited sample size also prevented us from assessing potential sex differences within and between the three groups. Finally, given the wide variability of clinical characterization in both ASD and SZ and more recent arguments in favor of subgrouping these heterogenous diagnostic entities into more homogenous sub-groups (e.g., Mottron & Bzdok, 2020; Oomen et al., 2022; Qi et al., 2020; Yan et al., 2021), acquiring larger samples is imperative for a thorough assessment of the relationship between INT and sensory processing in ASD and SZ.

In conclusion, the present study is a step forward in assessing the replicability of INT atypicalities in ASD and SZ. Despite current sample size limitations, it appears that the left lateral occipital gyrus and the right post-central gyrus hold the highest promise concerning replicable INT group differences between SZ and TD. Substantially larger samples are still required to definitely assess the robustness of the relationship between INT and psychopathology.

